# Cellular and mitochondrial effects of a gold-N Heterocyclic Carbene on LNCaP and PC3 prostate cancer cell lines

**DOI:** 10.1101/2025.01.25.634883

**Authors:** Judith Eguida Sangouard, Mathilde Cheron, Guillaume Rivrais, Serge Alziari, Federico Cisnetti, Arnaud Gautier, Isabelle Garreau-Balandier, Patrick Vernet

**Affiliations:** Université Clermont Auvergne, CNRS/IN2P3, LPC, F-63000 Clermont-Ferrand, France; Université Clermont Auvergne, CNRS, Sigma Clermont, ICCF, F-63000 Clermont-Ferrand, France

**Keywords:** Prostate cancer, Gold-N-heterocyclic carbene complex, Mitochondria, Anticancer agent

## Abstract

The discovery of cisplatin and its anti-tumor activities but also its side-effects led to the development of new cytotoxic metal complexes. Metal-based compounds and especially gold N-heterocyclic carbene complexes have shown their antitumoral activities and present a great interest of use in medicine. These metallodrugs have been shown to target nuclear DNA, but also other organelles such as mitochondria. The mechanisms of action have been studied and would involve the triggering of cell death, in particular by apoptosis, highlighting mitochondrial dysfunctions. The organometallic compound used in this study has been characterized by its IC_50_ and GI_50_ in two prostatic cancer cell lines (LNCaP and PC3). Its capacity to enter the cells and particularly mitochondria was also analyzed. Furthermore, its effects on several parameters (proliferation, cell death, cell cycle) have been determined in vitro. The compound displayed a cytotoxic effect (IC_50_ < 5µM) confirmed by the MTT assays and cell death analysis by flow cytometry. It also displayed a cytostatic effect confirmed by the determination of the GI_50_, and induced a brief cell cycle arrest, depending on the cell line and the metallocarbene concentration. Furthermore, the incubation with the gold N-heterocyclic carbene led to mitochondrial membrane potential changes and decreased the accumulation of the mitochondrial protein Rieske. All our results show a global cytotoxic and cytostatic effects of the gold N-heterocyclic carbene that can involve mitochondria.

## Introduction

Cancers are the second most common cause of death in the world. All over the world, prostate cancer has an annual incidence of 1.4 million and causes 375,000 deaths, making it the second most common cancer in men and the fourth most common cancer overall (« Prostate Cancer Statistics | World Cancer Research Fund International », s. d.). This mortality rate tends to decrease these last years, thanks to the democratization of screening tests allowing its early detection. However, prostate cancer remains a public health issue because the therapies are not always effective. Several therapeutic approaches are used, *ie.* radiotherapy, surgery and chemotherapy, alone or in combination. Chemotherapy is a systemic treatment that is the most often used on metastatic or resistant cancers. It is based on platinum(II)-based drugs such as cisplatin, carboplatin, oxaliplatin, nedaplatin, lobaplatin and heptaplatin. The mechanism of action is mediated through the platinum(II) crosslink with guanine residues, which induces DNA strand bending, DNA replication and transcription arrest or an interference with DNA repair, leading to cell growth and proliferation arrest through apoptosis (Dasari & Bernard Tchounwou, 2014; Jamieson & Lippard, 1999; Jung & Lippard, 2007; Pages et al., 2017). However, nowadays, its use is limited to few medical conditions due to severe side effects (nephrotoxicity, ototoxicity, neurotoxicity), non-selectivity, intrinsic and acquired chemo-resistance. This has prompted the development and research of new metal-based drugs (Becker et al., 2024; Cisnetti & Gautier, 2013; Gong et al., 2024). Regarding the metals, gold (auranofin, myocrisin and solganol), ruthenium (NAMI-A and KP1019) and copper (Casiopeinas) are mostly used in cancer therapies (Asif et al., 2016; Colina-Vegas et al., 2016; Leijen et al., 2015; Roder & Thomson, 2015).

Among the organometallic compound candidates, metal N-heterocyclic carbene complexes have emerged and represent a growing field of research (Liu & Gust, 2016; Van Der Westhuizen et al., 2021; Zhao et al., 2024). Various types of them exist with numerous metal centers and various functionalization of the carbene ligands (differing in ring size, nature of bond within the ring and the presence and nature of heteroatoms). Overall, both metal and ligand confer them different chemical and biological properties (Dahm et al., 2015; Haque et al., 2015; Verlinden et al., 2015; Zhang et al., 2018). Their reactivity towards specific targets and the chemical stability of the metal-carbene bond allow them to be used in human health (Scott & Nolan, 2005) and gold-carbene complexes have been demonstrated to possess a high stability under physiological conditions (Baker et al., 2005; Gautier & Cisnetti, 2012; Hickey et al., 2008; Holenya et al., 2014; Pratesi, 2016). Several metal N-heterocyclic carbene complexes based on silver, platinum or gold have demonstrated their anti-tumor properties *in vitro* and *in vivo* (J. Jiang et al., 2023; Y. Li et al., 2014; Wan et al., 2021).

Moreover, these compounds can specifically accumulate in tumor sites via Enhanced Permeability and Retention effect (EPR) (Kang et al., 2019; Oehninger et al., 2013) or by the use of ligands that recognize specific receptors. Some compounds have been reported to preferably target the nucleus or mitochondria (and more precisely DNA), or proteins like glutathione reductase (GR) and thioredoxin reductase (TrxR), key enzymes in the cells redox balance maintenance (Hickey et al., 2008; McCartin et al., 2022; Urig et al., 2006; Visbal et al., 2016). Due to their fundamental and central role in the cell, mitochondria might be an interesting target. Indeed, the implication of these dynamic organelles in cell survival, motility and death, metabolic reprograming that occur during cancer initiation and progression are well known. These events are characterized as cancer hallmarks (Hanahan & Weinberg, 2011; Z. Jiang et al., 2024). Moreover, mitochondria are the center of energy metabolism and are also involved in apoptosis, reactive oxygen species production, making them crucial for the fate of the cell (Dhingra & Kirshenbaum, 2014). Thus, several mitochondrial-targeting cytotoxic metal complexes have been synthetized and studied (Erxleben, 2019; Olelewe & Awuah, 2023). The anti-tumor properties of metal complexes of N-heterocyclic carbenes would involve the induction of mitochondrial cell death, inhibition of thioredoxin reductase (TrxR), estrogen receptor (ER) and cyclooxygenase (COX) in cases of breast cancer, lung cancer, leukemia, glioblastoma, prostate cancer, colon cancer (Barnard & Berners-Price, 2007; Eloy et al., 2012; Fan et al., 2019; Gautier & Cisnetti, 2012; Hickey et al., 2008; Holenya et al., 2014; Leitão et al., 2023; Y. Li et al., 2014; Liu & Gust, 2012; Pratesi, 2016; Salmain et al., 2023; Scalcon et al., 2018).

In our study, we explored the antitumor effects of the gold(I)-carbene complex **I** (**Fig. 1**) on two prostate carcinoma cell lines: LNCaP (PSA-expressing prostate cancer cell line) and PC3 (non-PSA-expressing prostate cancer cell line). Besides their PSA status differences, these two cell lines also differ from their AR, PTEN and P53 status (Cunningham & You, 2015; Horoszewicz et al., 1980; Kaighn et al., 1979). On the two cell lines, the internalization of **I** was analyzed, and its cytotoxicity was determined through several assays. In order to clarify its mechanism of action, the cell cycle and mitochondrial functions were also investigated. And finally, we also sought to highlight the type of cell death involved. This study gives a new insight of the antitumor effects of this gold-carbene against two different prostate cancer cell lines.

**Fig. 1.**
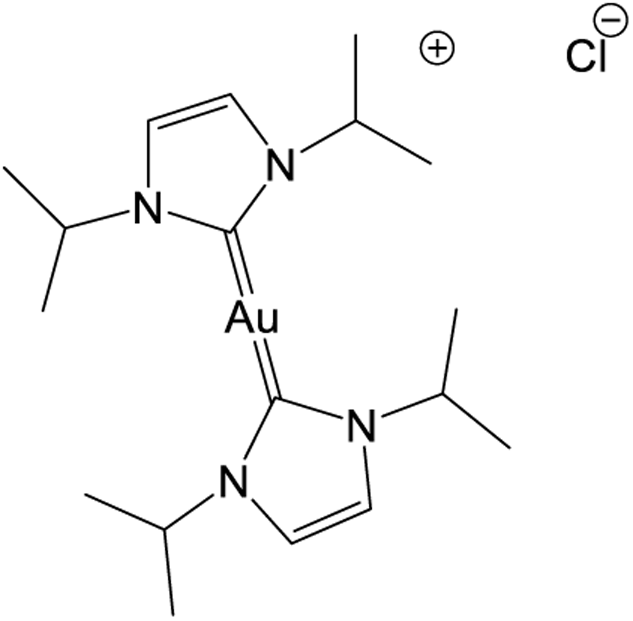
Chemical structure of the gold N-heterocyclic carbene **I** bis(N,N-diisopropylimidazol-2-ylidene)gold(I) chloride

## Material and Methods

### Au (I) N-Heterocyclic Carbene (NHC)

The gold N-heterocyclic carbene **I** was synthetized as described in the literature (Hickey et al., 2008). For the experimentations, different concentrations were obtained from a stock solution of **I** and dilutions were prepared in dimethyl sulfoxide (DMSO - Sigma).

### Cell culture

Human prostate cancer cells LNCaP (lymph node-metastasized cells) and PC3 (bone-metastasized cells) were purchased from the American Type Culture Collection (ATCC) and cultured in RPMI 1640 (Gibco by Life Technologies) supplemented with 10% heat inactivated fetal calf serum (tetracycline free, Biowest) and 0.2% gentamicin (Gibco by Life Technologies). The cells were maintained at 37°C in a humidified atmosphere of 5% carbon dioxide.

### Cells treatment

The cells were seeded and treated with various defined concentrations of **I**. They were pre-incubated for 3 hours followed by 0-120 hours incubation. Several controls were performed: cells with no treatment and cells treated with 0.01% of DMSO which is the concentration of DMSO in the wells treated with **I**, regardless of its final concentration.

### Determination of IC_50_ and cell proliferation assay

3-(4,5-dimethylthiazol-2-yl)-2,5-dyphenyltetrazolium bromide (MTT) assays were used to assess the cytotoxicity of the compound. First, the half maximum inhibitory concentration (IC_50_) of the compound was determined. The cells were briefly seeded in 96-well plates with 200 µL medium and then treated with different concentrations of the compound ranging from 10^-9^ M to 2.10^-5^ M for 48 hours. After treatment, the medium was discarded, then 100 µL of medium and 20 µL of MTT were added for 4 hours. The purple formazan crystals were dissolved by adding 100µL of 10% Triton-X 100 in acidic isopropanol (0.1 N HCl) and shaking the plates for 30 minutes. The absorbance was measured at 570 nm using a microplate reader. All conditions were tested in triplicate and repeated three times. The IC_50_ value was calculated from a dose response curve.

Secondly, the effects of **I** on survival were assessed. For that, the cells were seeded and then treated with different concentrations of the gold-NHC complex **I** for 0-120 hours. MTT assays were performed daily as described above.

### GI_50_ determination

In order to analyze the cytostatic activity of the gold-NHC complex **I**, the 50% growth inhibition (GI_50_) was determined by a clonogenic assay. The cells were seeded for 24 hours at a low density (500 cells/well), using three experimental parallels. They were then treated with various concentrations of the gold-carbene ranging from 0 to 4.10^-6^ M. Fourteen days later, the cell colonies were fixed with methanol at 37°C (Sigma-Aldrich) for 15 minutes, then stained with a 0.1% crystal violet solution (Sigma-Aldrich) and finally rinsed with water in order to remove excess staining. Only colonies of at least 50 cells were counted and the GI_50_ value was determined from the dose-response curve obtained.

The impacts of short time exposure with **I** instead of 14 days incubation were also determined. For that, the medium was changed either 3 or 24 hours after treatment; the cells were then incubated in a medium without the gold-NHC complex **I** and the experiment was proceeded 14 days later as described above.

### Cellular and subcellular gold intake

The quantity of gold internalized within the cells was determined through Inductively Coupled Plasma Mass Spectrometry (ICP-MS; Agilent 7500 Series). ICP-MS is a versatile method used to quantify non-endogenous metals in complex mediums and cells (Pröfrock & Prange, 2012).

The cells were cultured in 10 cm Petri dishes. After an incubation with **I** (0.5 µM for 3 hours), the medium was discarded and the wells were rapidly washed twice with phosphate buffered saline (PBS) and the cells were collected, digested with concentrated acids (HCl 12.4M, HNO_3_ 15.8M) in closed Savillex beakers in order to dissolve all organic and inorganic structures. The solutions were then evaporated at 100°C and afterwards cooled. Each dried sample was solubilized once again in a solution (HNO_3_ 0.4M, HF 0.05M) which contained 2 ppb ^115^In. This element was used as an internal standard to monitor and correct the sensitivity drift of the ICP-MS if necessary. The beakers were closed and placed at 90°C overnight. Once cooled, they were placed in an ultrasonic bath during 1 hour and after that, if necessary, some samples were diluted with the same previous ^115^In solution in order to obtain Au concentrations ranging from 1 ppb to 2.5 ppb.

The analysis were then performed in Argon plasma mode (1550W). The measured gold concentrations were systematically corrected for blank contribution; the instrumental drift, when significant, was used to correct the Au concentrations. Finally, the quantity of gold was expressed relatively to the number of cells.

In order to assess the quantity of gold in subcellular compartments, the cells were cultured and treated as described above and a mitochondria fractionation kit (Abcam) was used to separate the cytosol, mitochondria and nucleus according to the manufacturer’s instructions. The purity of the different subcellular fractions was controlled by western blot (using H3, Rieske and PDH, GAPDH antibodies respectively for the nucleus, mitochondria and cytosol) and they were then submitted to ICP-MS. The blotting protocol is described below. The percentage of gold in every fraction was calculated in comparison with total quantity of gold in the cells.

### Annexin V / Propidium Iodide staining

Flow cytometry studies in conjunction with annexin-V staining were carried out using the Dead Cell Apoptosis Kit with Annexin V Alexa Fluor 488 & Propidium Iodide (PI) (Molecular Probes) according to the manufacturer’s instructions. The cells were treated as described in the cells treatment paragraph. Positive controls (apoptosis induction) were produced by incubation with 2 μM staurosporine for 4 hours.

Adherent and floating cells were subsequently subjected to flow cytometry (FACS Calibur BD Biosciences; CYSTEM UCA PARTNER Platform) after being collected, washed and stained with fluorescein-labelled annexin-V (determined using a 488 nm blue laser and a 530/30 nm filter) and PI (determined using a 488 nm blue laser and a 575/24 nm filter). On average, 10000 events were analyzed in triplicate for each sample. A process of fluorescence compensation was used to correct the spectral overlap between Alexa Fluor 488 and PI. The data were analyzed with Flowing Software (Cell Imaging Core, Turku Centre for Biotechnology).

### Cell cycle analysis

The cells were seeded prior to the incubation with **I** at three different concentrations and were either collected after 3+ (0, 24, 48, 72 or 96h). The cell cycle was analyzed by flow cytometry (FACS Calibur BD Biosciences; CYSTEM UCA PARTNER Platform). For that, the wells were washed twice with PBS and the cells were collected, washed again and fixed with ethanol 70% at -20°C and kept for at least a day at -20°C. Further, they were washed and resuspended in PBS for 30 minutes at 37°C and 5.10^5^ cells were resuspended in 250µL of FxCycle PI/RNase Staining solution (Molecular Probes) for 30 minutes in the dark. The fluorescence intensity of 3.10^4^ events was measured using a 488 nm laser and a 575/24 nm filter. The data were analyzed with ModFit software (Modfit LT, The Verity Team).

### Preparation of cell lysate and western blot analysis

The cells were cultured and treated as described above. After the incubation with **I**, they were harvested and disrupted by sonication in Laemmli buffer and the proteins (20-50µg) were separated by SDS-PAGE and transferred onto a nitrocellulose membrane which was afterwards probed with relevant primary antibodies and HRP-conjugated secondary antibodies. The proteins were detected using Luminata forte western HRP Substrate (Millipore) or SuperSignal West Femto Maximum Sensitivity Substrate (Thermo Scientific) and visualized with a Fusion FX (Vilber Lourmat). The relative abundances of various candidate proteins were evaluated through a densitometric quantification of the signal intensity using Fusion FX Software (Vilber Lourmat). The following antibodies were used: β-tubulin, LC3 and Rieske.

### Mitochondrial membrane potential

The measurement of the mitochondrial membrane potential (ΔΨm) was carried out on approximately 5.10^5^ cells using 5.5’,6.6’-tetrachloro-1,1’,3,3’-tetraethylbenzimidazolycarbocyanine iodide (JC-1, Sigma Aldrich). JC-1 is a lipophilic, cationic dye that accumulates in active mitochondria under the form of aggregates (red fluorescence, 590nm) or monomers (green fluorescence, 529nm) during the fall of ΔΨm (Elefantova et al., 2018). After different incubation times in presence of **I**, the cells were collected and then incubated at 37°C for 20 minutes in the presence of the JC-1 probe. The cells were therefore analyzed by flow cytometry using two channels: FL1 (488 laser and 530/30 filter) and FL2 (488nm laser and 585/42nm filter). In average 3.10^4^ events were analyzed per sample. Cells treated with CCCP (100µM) for 20 minutes at 37°C served as a positive control. The data was analyzed using BD CellQuest Pro © software (Becton, Dickinson and Company).

### Statistical analysis

All experiments were performed independently at least in triplicate. The results are expressed as the mean value ± SEM. For statistical analysis, Xlstat software (Addinsoft) or GraphPad Prism (Graph PAD, San Diego, USA) was used. Data was analyzed using the Kruskall-Wallis test followed or not by Bonferroni correction. Pairwise multiple comparisons were analyzed by Dunn’s or Conover-Iman’s tests. Statistical significance was set at p < 0.05.

## Results and discussion

### Cytotoxic activity

Gold compounds have been investigated for their antitumoral properties and for their stability under physiological conditions (Bertrand et al., 2015; Dabiri et al., 2019; Roder & Thomson, 2015; Zhang et al., 2018). The gold-NHC complex used in our studies showed a cytotoxic activity against the breast cancer cell line MDA-MB-231 (Hickey et al., 2008). Here, we analyzed its cytotoxicity using two prostatic cell lines PC3 and LNCaP which respectively displayed IC_50_ at 4 µM and 2.4 µM (**Table 1**). These two values are of the same order (µM) but we can deduce that LNCaP cells seem slightly more sensitive to the gold-NHC **I** compared to PC3 cells. In addition, these values can be compared to IC_50_ obtained in studies where gold-carbene complexes effects have been studied against prostate cancer cells (Curado et al., 2019; Mui et al., 2016; Walther et al., 2020; Zhang et al., 2018). There is also the example of auranofin, an organometallic compound containing gold, used for the treatment of rheumatoid arthritis but also used for its anti-cancer characteristics. In their study, Zhang and colleagues showed that the cytotoxicity of this compound, through the inhibition of mitochondrial activity and the induction of oxidative stress, would be a consequence of its binding to TrxR (Zhang et al., 2019).

**Table 1.** IC_50_ values of the gold-NHC complex **I** in LNCaP and PC3 cell lines.

| Cell lines | IC<sub>50</sub> (μM) <sup>(a)</sup> |
| --- | --- |
| LNCaP | 2.4 |
| PC3 | 4 |

(a): The results are determined with a dose-response curve using the mean of at least three independent experiments.

The differences between the two cell lines can be linked to their PSA, AR, PTEN and/or P53 status (Cunningham & You, 2015; Horoszewicz et al., 1980; Kaighn et al., 1979). Indeed, prostate epithelial cells proliferation and differentiation depend on AR signaling. Additionally, the expression of PSA, used as a marker of prostate cancer cells growth, depends on AR signaling (Auchus & Sharifi, 2020; Desai et al., 2022). P53 is a transcription factor that plays a pivotal role in regulating the cell cycle and the cell’s ability to repair DNA damage (Chappell et al., 2011) and PTEN is a tumor suppressor gene. In addition, the loss of P53 activity modifies the specificity of AR to regulate gene expression in the context of prostate cancer (Guseva et al., 2012).

PC3 cells do not express AR nor PSA, unlike LNCaP cells which proliferation depends on androgens (Horoszewicz et al., 1980; Kaighn et al., 1979). A study of the effects of **I** on the expression of PSA and AR could support this hypothesis. Furthermore, PC3 cells express neither P53 nor PTEN unlike LNCaP cells which only express p53, making them potentially more sensitive to physicochemical and biological damages due to the presence of **I**.

### Cytostatic activity (GI_50_)

The analysis of the cytotoxicity of **I** showed its capacity to inhibit the two cell lines metabolic activities and therefore its potential to inhibit cell growth. So, the effects of the compound on cell growth have been analyzed by determining its growth inhibition (GI_50_) using a clonogenic assay. GI_50_ is the concentration of the gold-carbene necessary for the inhibition of 50% of cell growth. In our cell culture conditions, due to the inability of LNCaP cells to clone (cloning efficiency of 4% (data not shown)), (Bennett et al., 1997) this experiment was only carried out on PC3 cell line.

The cells were seeded at low density and treated with different concentrations of **I**. The colonies with at least 50 cells were counted after a crystal violet staining 14 days later. Three different experimental conditions were developed: the cells were in contact with the metallocarbene either for 14 days or only for 24 or 3 hours. The results are presented in **Table 2**.

**Table 2.** Analysis of the effect of the carbene on cell growth using a clonogenic assay. The percentage of survival is expressed in relation to the “DMSO” condition ± SEM. The results are from at least 5 independent repetitions.

| (I) | 0μM | 0.5 μM | 1μM | 2μM | 4μM |
| --- | --- | --- | --- | --- | --- |
| 3h | 100 | 93.0 ± 5.4 | 94.4 ± 11.3 | 80.6 ± 16.8 | 73.7 ± 11.1 |
| 24h | 100 | 93.5 ± 3.7 | 81.2 ± 5.9 | 35.2 ± 12.0 | 17.8 ± 6.6 |
| 14 days | 100 | 71.3 ± 9.4 | 28.4 ± 14.8 | 0 | 0 |

In the three conditions, **I** showed a dose effect and different profiles emerge depending on how long the metallocarbene has been in contact with the cells. Indeed, a weak cytostatic effect is observed when the gold-NHC is maintained in the medium for only 3 hours whatever its concentration. On the other hand, a strong cytostatic effect of **I** is noticed when it is maintained for the duration of the experiment (14 days). Unlike these two previous conditions, the concentration inhibiting half of the cell survival: GI_50_ = 1.7 µM had been estimated through an effect-dose curve (**Fig. 2**) with the condition where **I** is maintained for 24 hours.

**Fig. 2.**
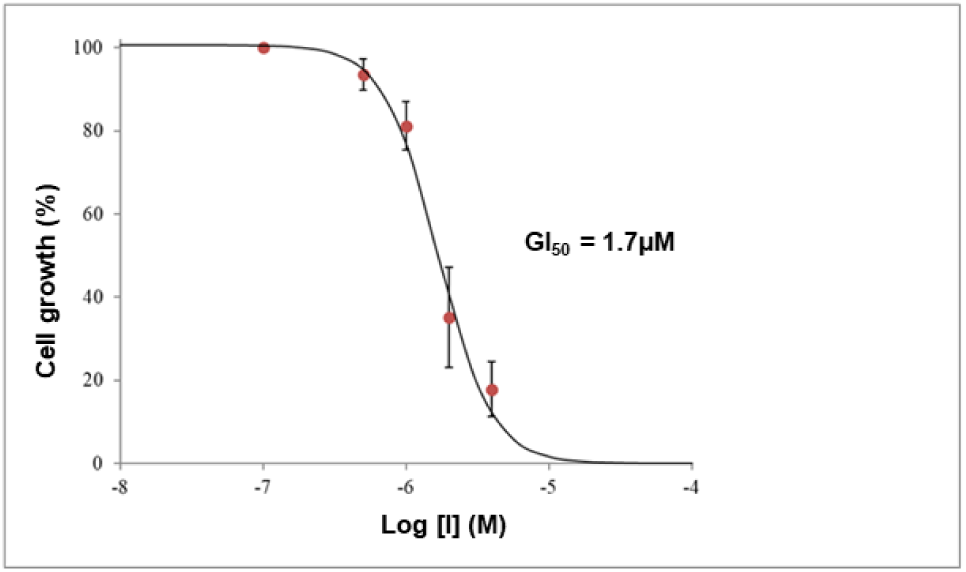
GI_50_ value of **I** in PC3 cell line. The carbene was maintained 24h in the medium and its effects on growth inhibition was observed 14 days later. Mean of at least 5 independent repetitions

These results show the impact of **I** on PC3 cells growth and the impact of maintaining **I** in the medium on cell division capacity: the longer the incubation time, the more quickly cell growth was impacted. The gold-NHC **I** has marked cytostatic capacities on PC3 cells. In addition, the values obtained are very similar to those found in the literature with other gold-carbenes on the PC3 cell line (Zhang et al., 2018).

### Cellular and subcellular uptake

The intracellular localization of organometallic compounds can influence and explain their mechanism of action. Some metal N-heterocyclic carbene complexes have been localized in the cell and more specifically in the cytoplasm and some showed a mitochondrial, lysosomal or endoplasmic reticulum preference (Jakob et al., 2020). As the mitochondrial localization of the metallocarbene used in our study has been demonstrated in the literature on MDA-MB 231 cell line (Hickey et al., 2008), we investigated this with LNCaP and PC3 cell models. First its internalization into the cells was analyzed by ICP-MS, then secondly, its subcellular distribution.

As shown in **Fig. 3A** (LNCaP) and **B** (PC3), both cell lines internalize gold since no signal was detected in the untreated cells while gold was quantified in treated cells. However, the accumulation of gold in the two cell lines seems to differ from one cell line to another but this could be explained by the difference in seeding and growing conditions between PC3 and LNCaP cell lines.

**Fig. 3.**
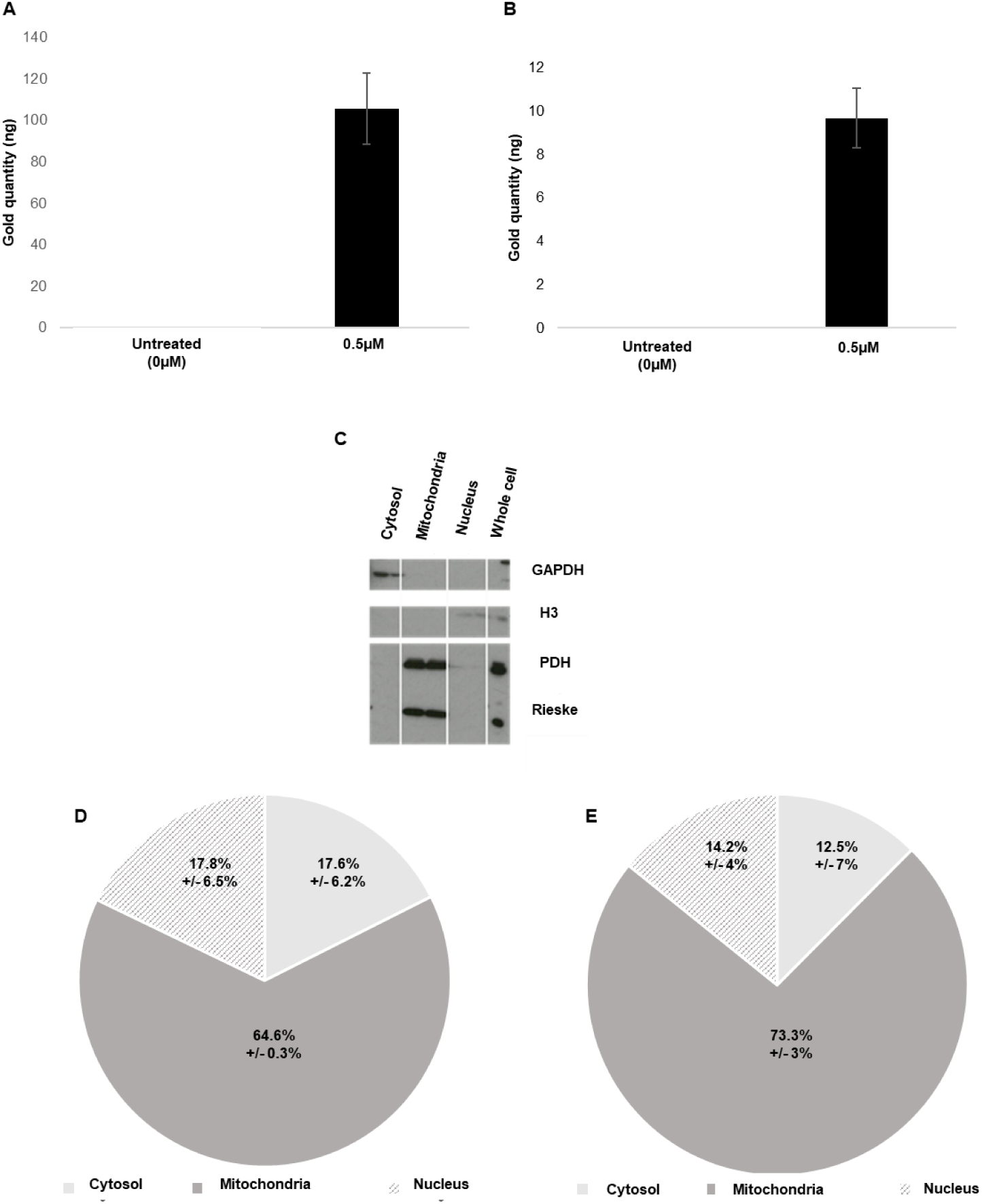
Gold uptake at cellular and subcellular levels by LNCaP (A, D) and PC3 (B, E) cells exposed to the gold-NHC complex **I** at 0.5 µM for 3 hours. The purity of the subcellular fractions was analyzed by western blot (C). Mean of three independent repetitions +/-SEM

Subsequently, cell fractionation was carried out on samples with the same incubation conditions (0.5 µM for 3 hours). Three compartments were isolated: nucleus, cytosol and mitochondria, validated by the purity of these fractions (**Fig. 3C**). The three fractions were subjected to ICP-MS and the distribution of gold in the 3 compartments was expressed as a percentage of the total quantity of gold in the cells. Gold is mainly found in mitochondria (64.58% ± 0.3% in LNCaP cell line and 73.33% ± 3% in PC3 cell line) (**Fig. 3D** and **E**). The proportion of gold inside the mitochondrial compartment is very similar from one experiment to another, regardless the cell type.

The observed mitochondrial enrichment is predictable given the charge and lipophilic properties of gold-NHC complexes evidenced in the literature (Hickey et al., 2008; Zhang et al., 2018). Additionally, an irreversible binding of metallocarbenes with selenoproteins such as mitochondrial thioredoxin reductase has been demonstrated in several studies (Holenya et al., 2014; Zhang et al., 2019; Zou et al., 2018). From a chemical point of view, this may be rationalized by considering that a softer donour such as a selenide is expected to bind more strongly to gold as compared to sulphide (Chaudiere & L. Tappel, 1984). Gold was also present in the cytosol and nucleus, in similar proportions. Its presence can be explained by a non-exclusive mitochondrial targeting. In addition, thioredoxin reductase also exists in cytosolic form that could also form a complex with gold-NHC complexes (Lu & Holmgren, 2014).

### Antiproliferative activity

The effects of **I** on cell proliferation was determined using the colorimetric MTT assay. Three different concentrations of **I** were analyzed at different times between 24 to 120h after a preincubation of 3 hours. The metallocarbene was maintained in the medium all along. Due to the low proliferative activities of the LNCaP cell line, only the PC3 cell line was studied.

The analysis of our data allow us to confirm that the presence of DMSO (diluted at 1/5000) has no effect on the PC3 cell line viability that is identical in both conditions (DMSO and medium) (data not shown). The DMSO condition was therefore used as the reference.

As shown in **Fig. 4**, the gold-NHC complex **I** displayed dose and time dependent antiproliferative activities against the prostate cancer cell line PC3. A concentration of 0.5 µM (8 times lower than the IC_50_) had very little effect on cell proliferation between 72 and 96 hours. On the other hand, the gold-NHC complex **I** at the concentrations of 2 and 4 µM had more significant impacts on cell proliferation from 48 hours, in a fairly similar manner, but a more pronounced effect at 96 and 120 hours at the concentration of 4 µM (IC_50_): at 2 µM the viability decreases from 69% ± 9% to 50% ± 7% between 48 and 120 hours compared to 63% ± 9% and 19% ± 1% at 4 µM. It could be explained by the hypothesis that at 2 µM cell death is induced but a large part of the cells retain their capacity to proliferate probably because of cell cycle arrest (the viability remains around 50% compared to the control), while at 4 µM cell death, or at least the inability to proliferate, would be prevailing since the viability shows a 2-fold decrease in 24 hours between 72 and 120 hours).

**Fig. 4.**
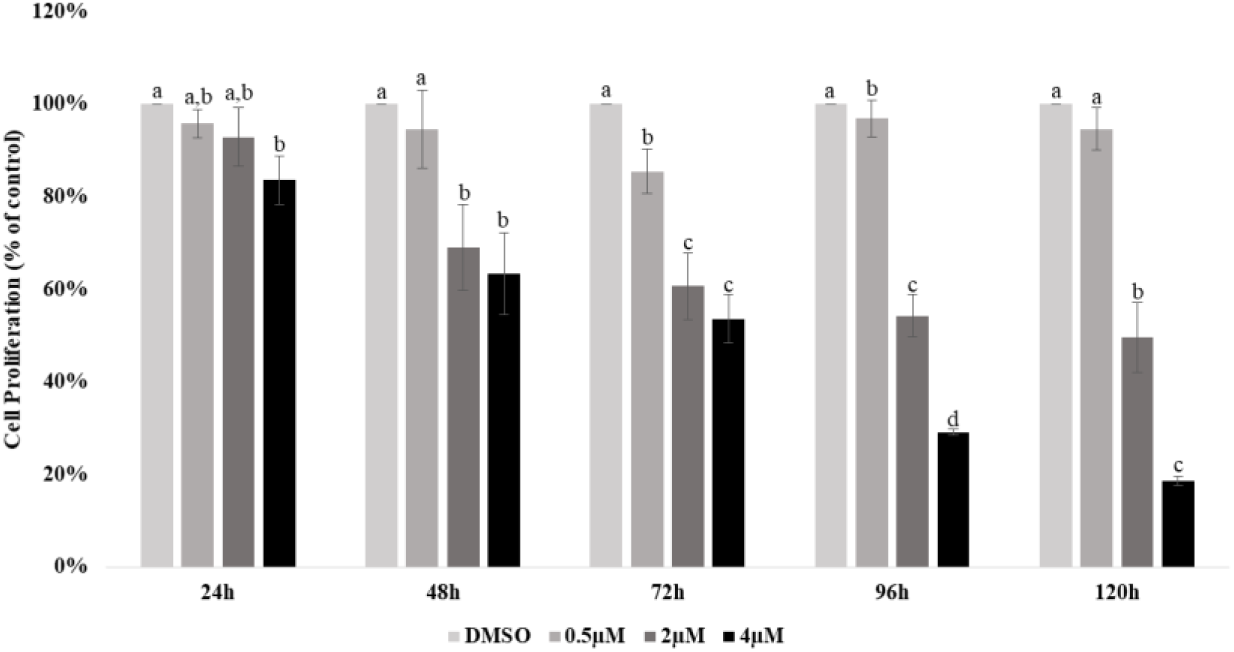
Antiproliferative capacities of the metallocarbene **I** on PC3 cells using the MTT colorimetric assay. The results are represented as mean+/-SEM of at least three independent experiments. Statistical comparisons are made for each time and significant differences (P < 0.05) are indicated by dissimilar superscripts

Beside gold, the antiproliferative capacities of metal N-heterocyclic carbene complexes have also been demonstrated with other metals such as silver(I), titanium(IV), platinum(II) and palladium(II) in cellular and mouse models of various types of cancers including prostate cancer. Those capacities are interesting from a therapeutic point of view because, compared to cisplatin, oxaliplatin or carboplatin, these organometallic compounds have superior or comparable antiproliferative capacities against cancer cells but weak antiproliferative capacities against healthy cells (Batnasan et al., 2024; Das et al., 2023; Jakob et al., 2020; Y. Li et al., 2014; Nicolás et al., 2024). Some of these studies demonstrated the activation of cell death pathways such as apoptosis, or cell cycle arrest.

### Induction of cell death

An analysis of apoptotic processes by flow cytometry, complemented by a western blot analysis of the accumulation of actors involved in autophagy (LC3) were carried out. Gold-NHC complexes have been shown to induce apoptosis via their irreversible binding to mitochondrial selenoproteins (Hickey et al., 2008; Schuh et al., 2012; Wang et al., 2011). Apoptosis is characterized by morphological modifications such as plasma membrane blebbing and nuclear fragmentation. Annexin V staining was used to detect phosphatidylserine externalization due to membrane blebbing and Propidium Iodide (PI) staining in order to label the nucleus of cells that have lost their plasma membrane integrity (features of necrosis). Thus, using these two markers, the analysis by flow cytometry allowed us to distinguish four cell populations: living cells (negative for both markers), cells in early apoptosis (positive for annexin V and negative for PI), cells in late apoptosis (double positive), and finally, those in necrosis (positive for PI and negative for annexin V) (Crowley et al., 2016). This distinction between late apoptosis and necrosis results from the fact that for some cell models, necrosis appears as a secondary step following the triggering of apoptosis (Honda et al., 2000).

The cells were seeded and treated at the defined concentrations of **I** for 0-96 hours. Staurosporine (STS) was used as a positive control. First, a significant difference is observed between the two cell lines regarding their response to STS treatment (**Fig. 5** and **6 / STS**). These behaviors described in the literature are due to the triggering of different cell death pathways (Marcelli et al., 2000).

**Fig. 5.**
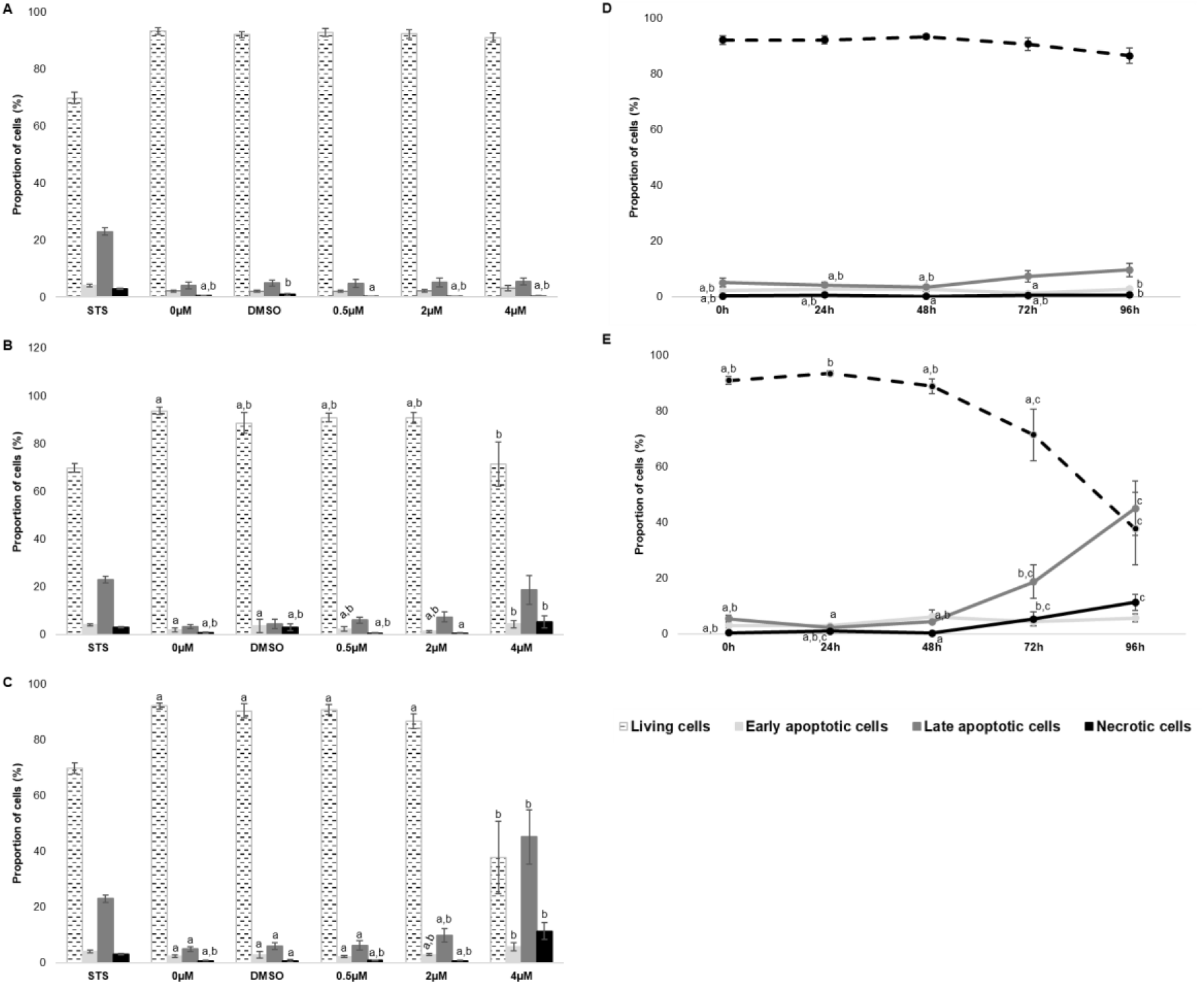
Analysis of apoptosis induction in PC3 cells treated with the gold-NHC complex **I**, at 0h (A), 72h (B) and 96h (C) via flow cytometry assays. The time effects of **I** at 2 µM and 4 µM are respectively represented in D and E. Staurosporine (STS) is used as a positive control. The results are the mean +/- SEM of at least three independent experimentations. Statistical comparisons are made for each cell population (living cells, early apoptotic cells, late apoptotic cells and necrotic cells). Means with dissimilar superscripts are significantly different (P < 0.05)

**Fig. 6.**
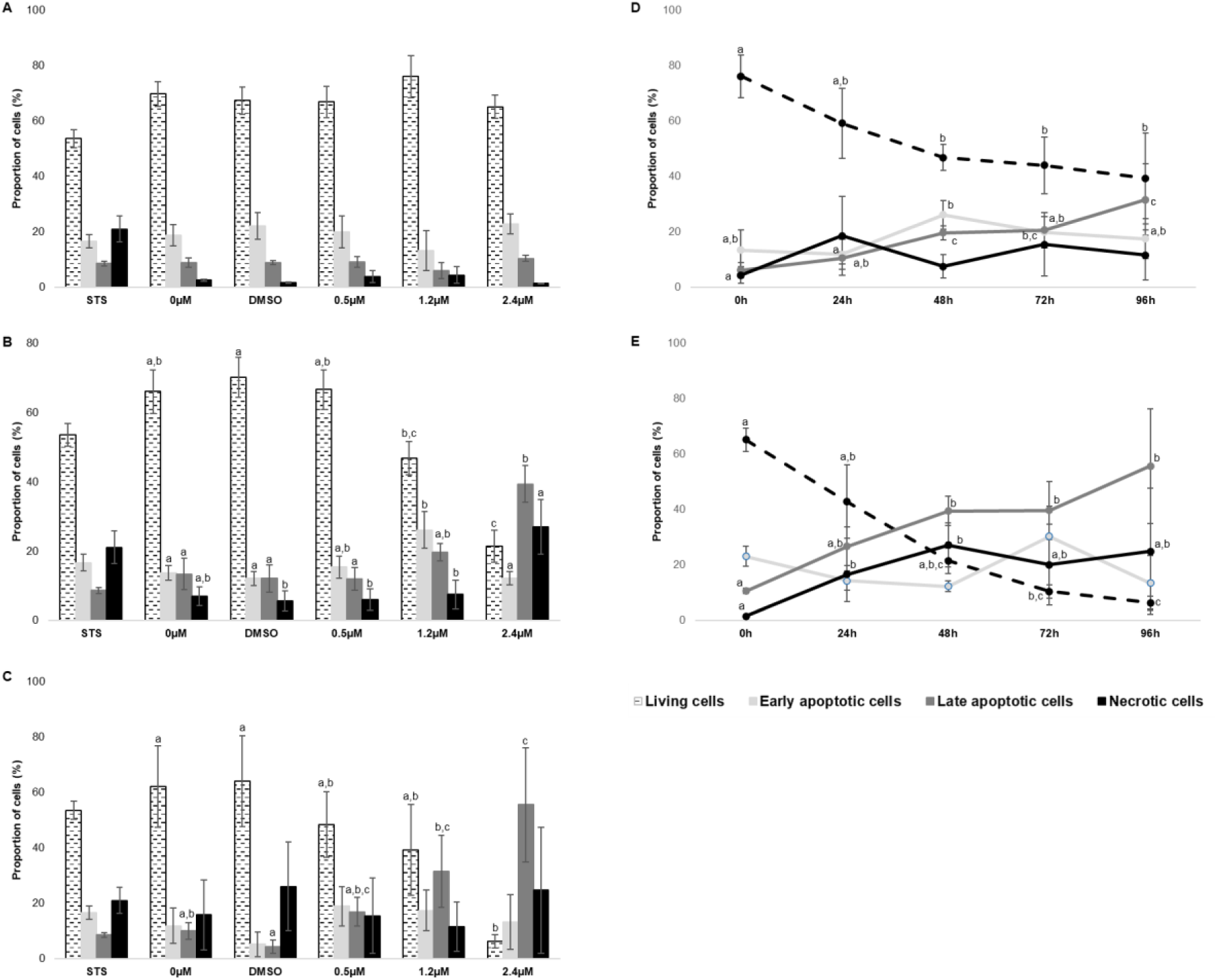
Analysis of apoptosis induction in LNCaP cells treated with the gold-NHC complex **I**, at 0h (A), 48h (B) and 96h (C) via flow cytometry assays. The time effects of the concentrations of 1.2 µM and 2.4 µM are respectively represented in D and E. Staurosporine (STS) is used as a positive control. The results are the mean +/- SEM of at least three independent experimentations. Statistical comparisons are made for each cell population (living cells, early apoptotic cells, late apoptotic cells and necrotic cells). Means with dissimilar superscripts are significantly different (P < 0.05)

The analysis of the distribution of PC3 cells in the different phases of apoptosis or necrosis (**Fig 5** and **SI. 1**) shows that the “0 µM” (medium only) and “DMSO” conditions present very similar profiles with approximately 95% non-apoptotic or necrotic cells. The gold-NHC **I** at 4 µM (IC_50_) induced an increase in the proportion of cells in early apoptosis at 72 hours followed by an increase in cells in early, late apoptosis and necrosis at 96 hours post treatment (**Fig. 5 A-C**). The observations made here, although surprising compared to cell proliferation test results (MTT assay) are not contradictory. Indeed, a significant drop in proliferation was observed in the presence of **I** at 4 µM but also at 2 µM 48 hours post treatment. This can be explained by the fact that the proliferation test is based on the detection of metabolic activities (Kumar et al., 2018) while the analysis carried out here involves other membrane-related processes. The results obtained at 24 h and 48h are presented in the supplementary information (**SI. 1**).

The LNCaP cell line showed a different overall behavior from the PC3 line, even for basal conditions. The analysis of the cells distribution in the different dead cells populations 48h post treatment (**Fig 6B**) showed an increase in the proportion of cells in early apoptosis for the 1.2 µM concentration in comparison to the DMSO condition (12% ± 2% (DMSO) vs 26% ± 5%). Also, an increase in the proportion of cells in late apoptosis is observed when the cells are treated with **I** at 1.2 µM and 2.4 µM (12% ± 4% vs 20% ± 2% for 1.2 µM and 12% ± 4% vs 39% ± 5% for 2.4 µM), associated to a decrease in the proportion of living cells (70% ± 6% vs 47% ± 5% and 70% ± 6% vs 21% ± 5% respectively for 1.2 and 2.4 µM); 96h post treatment, an increase in the proportion of cells in late apoptosis with a drastic decrease in living cells is observed with the same concentrations (**Fig. 6 A-C**). The results obtained at 24h and 72h are presented in the supplementary information (**SI. 2**).

It seems that the gold-NHC **I** would tend to engage PC3 cells towards an apoptotic pathway and for a small part of them (from 0 to 96h, only a slight increase from 0.3% to 0.74% and 0.4% to 11%), towards the necrotic pathway with the concentrations of 2 and 4 µM (**Fig. 5 D-E**). Concerning the LNCaP cell line, the cells seem quite sensitive to cell death, considering their behavior regarding the STS treatment and the evolution of the living cells populations over time and increased concentrations of the gold-NHC complex **I** and a significant proportion of cells with apoptotic and necrotic cells characteristics (**Fig. 6 D-E**). The differences between the two cell lines regarding the induction and kinetics of cell death would be linked to their P53 status (LNCaP cells express P53 contrary to PC3 cells) (Carroll et al., 1993). Indeed, P53 plays a central role in apoptosis initiation and also induces necrosis in response to DNA damages (Ying & Padanilam, 2016). The MTT assays and cytometry results would reflect a population of cells that are more or less metabolically active but still capable of maintaining survival despite the initiation of cell death process.

Several studies have reported the induction of apoptosis by metal-carbenes in various types of cancer cells (Holenya et al., 2014; McCall et al., 2017; Visbal et al., 2016). Other studies involving gold-NHC complexes also showed the activation of apoptosis through the inhibition of TrxR coupled with an increase in phospho-p53, which induces the cleavage and activation of caspase, a cleavage of PARP and inhibition of Bcl-2, an anti-apoptotic protein (Gupta et al., 2019; Holenya et al., 2014; Karaca et al., 2017; Nandy et al., 2016; Zou et al., 2018). Concerning prostate cancer more particularly, the induction of apoptosis was observed on PC3 cells following auranofin treatment at the concentration of IC_50_ after 72 hours (Curado et al., 2019), which is consistent with our data.

The characterization of the type of cell death involved was also carried out by western blot analysis of the cleavage and conversion of LC3, an actor involved in autophagy. LC3 exists in two forms: a free cytosolic form (LC3-I) and a form where the protein is associated with the phagosome (LC3-II) and whose electrophoretic mobility in SDS-PAGE is greater than that of the first form. Thus, an increase in the LC3-II/LC3-I ratio signifies an activation of autophagy (Giménez-Xavier et al., 2008).

The analysis were carried out on PC3 cell line as described (**Fig. 7**). Our results do not demonstrate any significant variation in the LC3-II/LC3-I ratio over time or in the presence of increasing concentrations of **I**. However, in their studies, Lee and colleagues demonstrated the activation of autophagy by NHC-like compounds associated with gold in cellular models of MCF-7 breast cancer cell line (Lee et al., 2017). To date, the study of the induction of autophagy by organometallic compounds in prostate cancer models does not exist. On the other hand, the induction of autophagy by gold nanoparticles is fairly well documented (J. J. Li et al., 2010). However, it must be kept in mind that the triggering of an autophagic process is not systematic in tumor progression since it can be activated or inhibited depending on the micro environmental conditions (Amaravadi et al., 2016; Mulcahy Levy et al., 2017; Ruan et al., 2019).

**Fig. 7.**
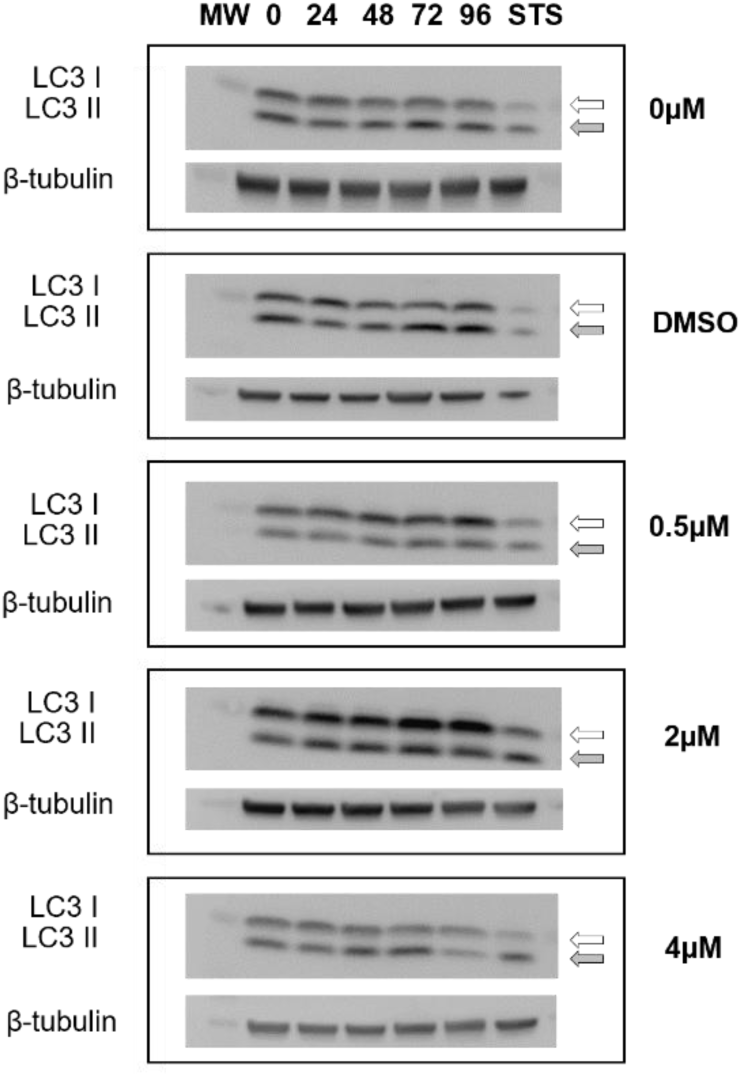
Effects of the gold-NHC complex **I** on the protein levels of LC3 in PC3 cells. Staurosporine (STS) was used as a positive control

### Cell cycle arrest

It has been demonstrated that metal N-heterocyclic carbene complexes induced cell cycle disruptions and thus cell cycle arrest, which is often associated with the cytotoxicity of those organometallic compounds (Ray et al., 2007; Teyssot et al., 2009). LNCaP and PC3 cells were seeded and treated with different concentrations of **I** (0.5 µM, IC_50_/2 and IC_50_) for the indicated times and cell cycle analysis was performed using PI staining and flow cytometry.

The analysis of the cell distribution in the cell cycle phases in the absence of treatment (0µM and DMSO) (**Fig. 8** and **SI. 3**) is in agreement with recent the data (Adekiya et al., 2024; Elagawany et al., 2024). The analysis of the cell cycle from 24 to 96 hours following the incubation with the gold-NHC complex **I** did not reveal a marked effect on PC3 cells (Supplementary information **SI. 3**). However, concerning the LNCaP cell line (**Fig. 8**), the most marked effect of the gold-NHC complex **I** is observed at 24h with a significant increase in the number of cells in G0/G1 associated with a decrease of the number of cells in phase S. These low variations do not appear to be dose-dependent. These disturbances are fleeting (24 hours and nothing beyond) and the brevity of the disturbances suggests an effect of the gold-NHC complex **I** on LNCaP cell cycle but also, in parallel, the capacity of these cells to quickly implement repairs. The possible arrest in G0/G1 phase with increase in subG1 phase is a strong signal to trigger apoptosis (Wu et al., 2016).

**Fig. 8.**
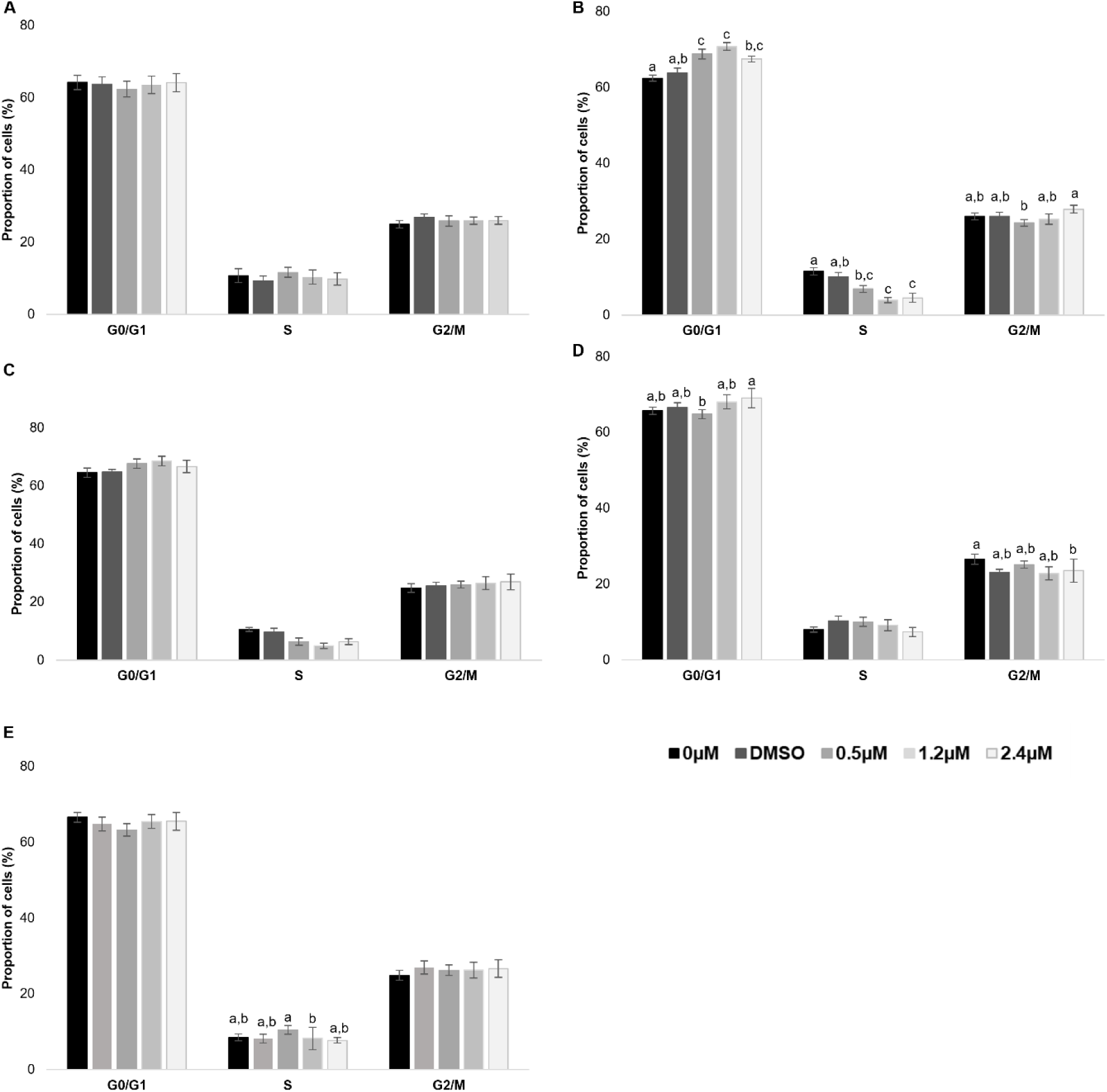
Distribution of LNCaP cells in the phases of the cell cycle after the treatment with the gold-NHC complex **I** at 0h (A), 24h (B), 48h (C), 72h (D), 96h (E).The results are represented as the mean of at least three independent cell cycle analysis. . Statistical comparisons are made for each phase of the cell cycle. Means with dissimilar superscripts are significantly different (P < 0.05)

### Mitochondrial disturbances

An important step in the apoptotic process is the mitochondrial membranes permeabilization, which leads to the release of pro-apoptotic proteins into the cytosol. More generally, the loss of mitochondrial membrane potential is a particularly reliable predictor of apoptosis. It has been reported that cationic complexes can pass through the mitochondria lipid membrane and induce mitochondria-mediated-apoptosis (Berners-Price & Filipovska, 2011; Che & Sun, 2011; Hickey et al., 2008; Jellicoe et al., 2008). The membrane potential was studied by flow cytometry according to the protocol described. CCCP was used to disrupt mitochondrial potential and therefore as a positive control.

PC3 cells showed a dose-dependent mitochondrial membrane depolarization from 24h with a significant effect of the gold-NHC complex **I** when the cells were treated with 4 µM (IC_50_). From 48 hours onwards, this effect was dose-dependent, but while it persisted at 4 µM, it faded over time for the lowest concentrations (**Fig. 9A**). Concerning the LNCaP cells (**Fig. 9B**), they displayed basal high values of depolarization (around 40%) and 3 hours post treatment. However, as for PC3 cells, the membrane potential is sensitive to the highest concentration of the gold-NHC complex **I** (IC_50_ - 2.4 µM) from 72 hours with, however, a more marked loss of membrane potential, equivalent to that obtained with CCCP, suggesting a greater sensitivity. Mitochondrial membrane depolarization has been demonstrated in several studies using metal N-heterocyclic carbene complexes associated with consequences on cell fate that depend on the cell type but also on the type of the metal N-heterocyclic carbene complexes used (Baker et al., 2006; Eloy et al., 2012; Li et al., 2015, 2020). We could logically think that a mitochondrial mediated apoptosis would be the consequence of the loss of mitochondrial membrane potential but our results might highlight a protection mechanism from cell death by apoptosis. Once again, the p53 status of PC3 cells could be involved.

**Fig. 9.**
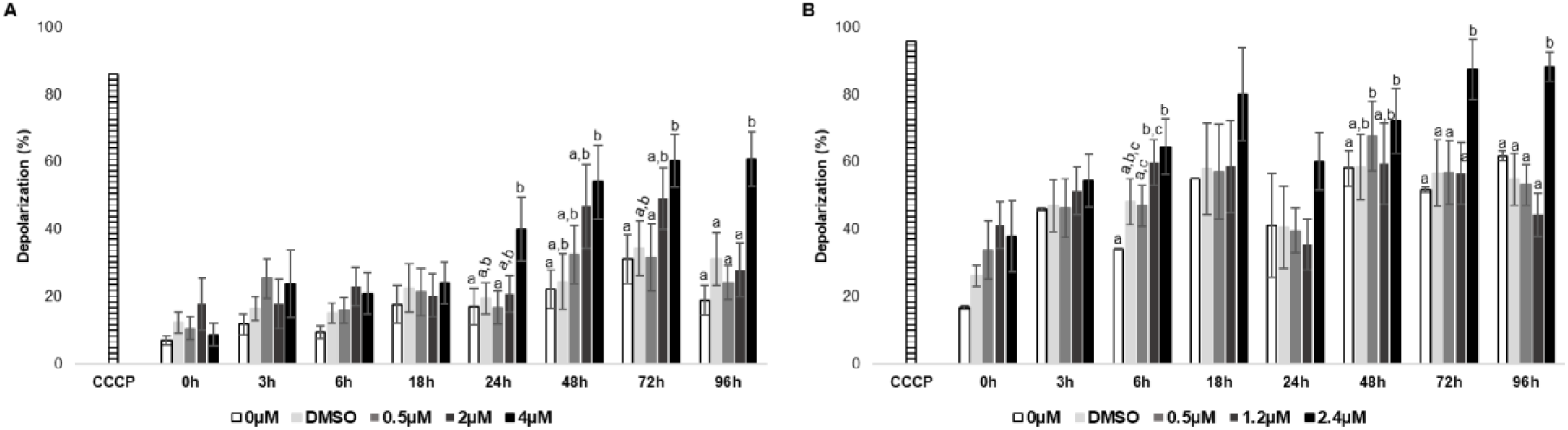
Effects of the gold-NHC **I** on the mitochondrial membrane polarization in PC3 (A) and LNCaP (B) cells assayed by flow cytometry. CCCP was used as a positive control and the results are the mean +/- SEM of at least three independent experiments. Statistical comparisons are made for each treatment time and means with dissimilar superscripts are significantly different (P < 0.05)

Several observations made previously on the localization of the gold-NHC complex **I**, cell survival, apoptosis, membrane potential, are significantly disturbed and globally tend to converge with disturbances of the mitochondrial metabolism. In order to estimate these changes, a western blot analysis of the accumulation of the protein Rieske, a subunit of the complex III was carried out.

The **Fig. 10** shows a stable quantity of Rieske over time when the cells were untreated (0µM) or treated with DMSO. When treated, a progressive disappearance of the protein in a time and dependent way was observed from 48h post treatment.

**Fig. 10.**
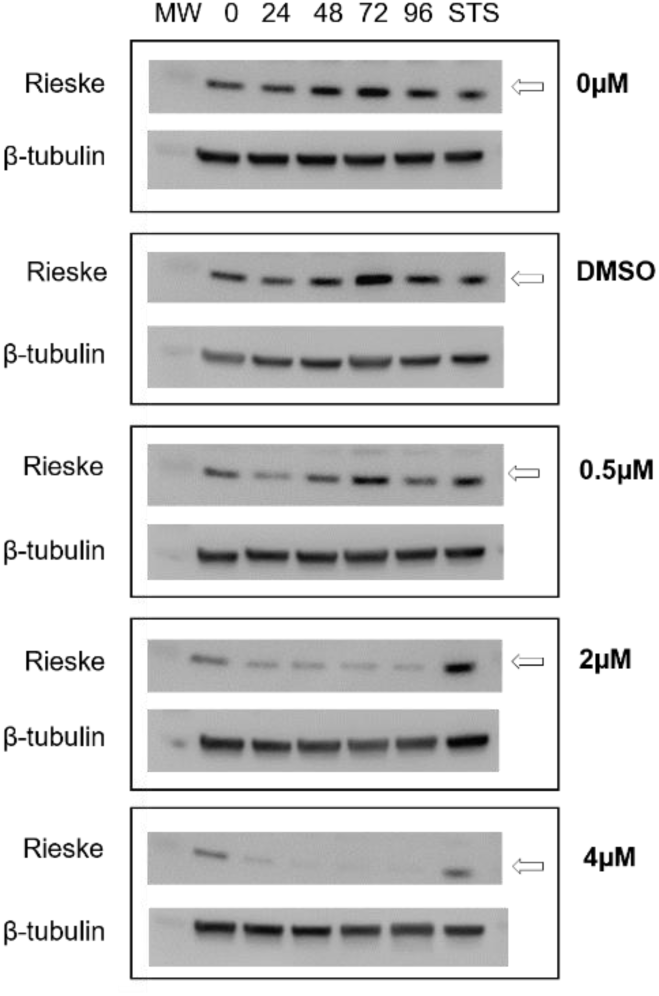
Effects of the gold-NHC **I** on the protein level of Rieske in PC3 cells. Staurosporine (STS) was used as positive control

The Supplementary information **SI. 4** illustrates the result obtained when the cells are in the presence of the gold-NHC complex **I** at the concentration of 4 µM for only 3 hours. The gold-NHC complex **I** was then eliminated by changing the medium. Interestingly, the low accumulation of the protein from 0h to 24h is reversed and gradually increased after until 96h. This could indicate that **I** may play a role during the synthesis of the protein and that this effect is reversible since the removal of the gold-NHC complex **I** in the medium allowed back the accumulation of the protein. The protein Rieske has a 2Fe-2S center in its Fe^III^ state, conducive to exchanges with gold(I), which could explain our results (Ali et al., 2014; Ouyang et al., 2018). These experimentations are only made once and need to be confirmed.

## Conclusion

In conclusion, we have characterized the actions of the gold-NHC complex **I** on two prostate cancer cell lines (LNCaP and PC3). **I** has been used previously against breast cancer cells (Hickey et al., 2008). Several cellular and mitochondrial aspects were analyzed in our studies. First of all, our results confirmed the mitochondrial localization of the metallocarbene in the two cell lines which for the rest of the experiments showed different behaviors that could be linked to their different PSA, P53, PTEN, AR statuses. Our biological studies allowed to highlight its cytotoxic and cytostatic capacities: in fact, the gold-NHC complex **I** inhibited cell proliferation and growth. This was manifested by a slight increase in apoptotic and necrotic populations (only at high concentrations) in PC3 cells but more marked in LNCaP cells. No effect on autophagy was observed. Cell cycles did not seem to be impacted (apart from a brief arrest in G0/G1 phase for the LNCaP line), but a loss of mitochondrial membrane potential and also the expression of the protein Rieske, a complex III protein were observed. These observations are promising regarding the development of drugs targeting mitochondria. Further tests are required in order to determine the molecular mechanisms involved, the importance of AR and P53 signaling and it would be also interesting to analyze the effects of the compound on other prostate cancer cell types. Finally, given its characteristics, it will also be interesting to study the effects of this gold-NHC complex coupled to radiotherapy on prostate cancer cells.

## Acknowledgement

The authors would like to thank Maud BOYET and Jean-Luc PIRO (*Laboratoire Magmas et Volcans*) for allowing us to carry out the ICP-MS experiments.

This work was supported by the MESR; Clermont Auvergne University; the LABEX PRIMES (ANR-11-LABX-0063) of Lyon University, within the program “*Investissements d’Avenir*” (ANR-11-IDEX-0007) operated by the French National Research Agency (ANR). The funders had no role in the study design, data collection and analysis, decision to publish, or preparation of the manuscript.

## Statements and Declarations

Authors do not have any financial or non-financial interests that are directly or indirectly related to this work.

## Authors’ contributions

Alziari S., Garreau-Balandier I., Cisnetti F., Eguida Sangouard J., Gautier A., and Vernet P. contributed to the study conception and design. Material preparation, data collection and analysis were performed by Eguida Sangouard J., Cheron M., Garreau-Balandier I. and Rivrais G. .The first draft of the manuscript was written by Eguida Sangouard J. . Garreau-Balandier I., Cisnetti F., Eguida Sangouard J., Gautier A., and Vernet P. contributed to editing the manuscript. All authors read and approved the final manuscript.

