## Supplementary information for "Cellular and mitochondrial effects of a gold-N Heterocyclic Carbene on LNCaP and PC3 prostate cancer cell lines"

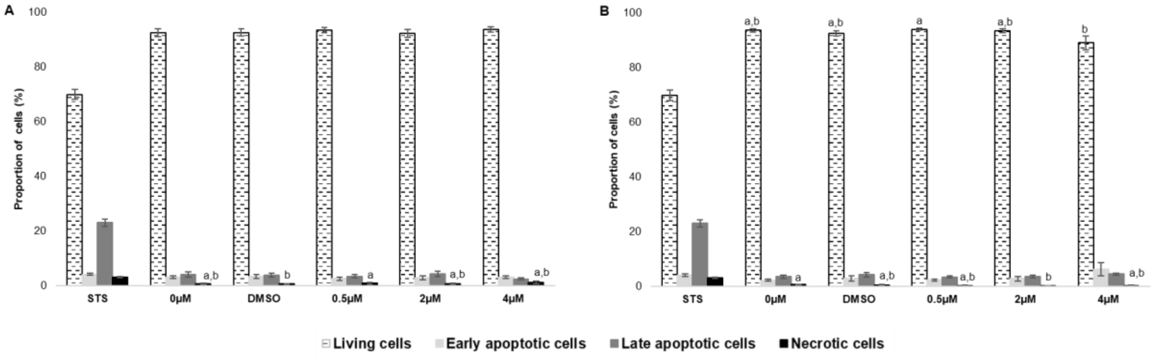


**SI. 1** Analysis of apoptosis induction in PC3 cells treated with the gold-NHC complex **I**, at 24h (A) and 48h (B) via flow cytometric assays. Staurosporine (STS) was used as a positive control. The results are the mean +/- SEM of at least three independent experimentations. Statistical comparisons are made for each cell population (living cells, early apoptotic cells, late apoptotic cells and necrotic cells). Means with dissimilar superscripts are significantly different ($P < 0.05$)

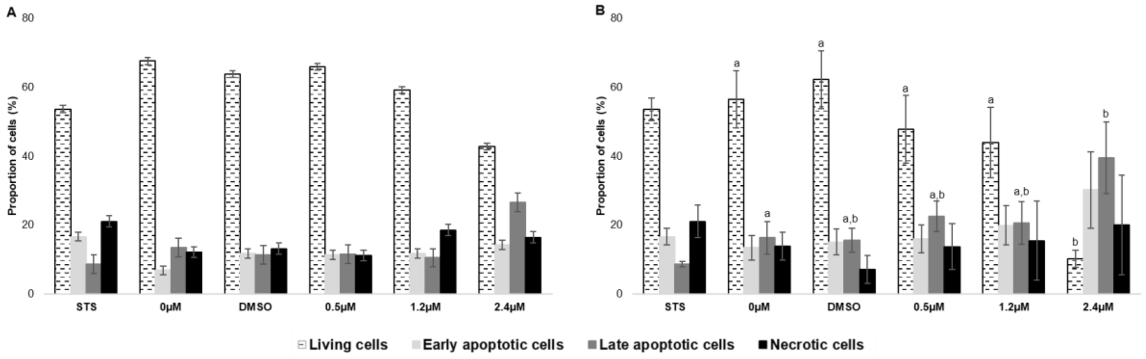


**SI. 2** Analysis of apoptosis induction in LNCaP cells treated with the gold-NHC complex **I**, at 24h (A) and 72h (B) via flow cytometric assays. Staurosporine (STS) is used as a positive control. The results are the mean +/- SEM of at least three independent experimentations. Statistical comparisons are made for each cell population (living cells, early apoptotic cells, late apoptotic cells and necrotic cells). Means with dissimilar superscripts are significantly different ($P < 0.05$)

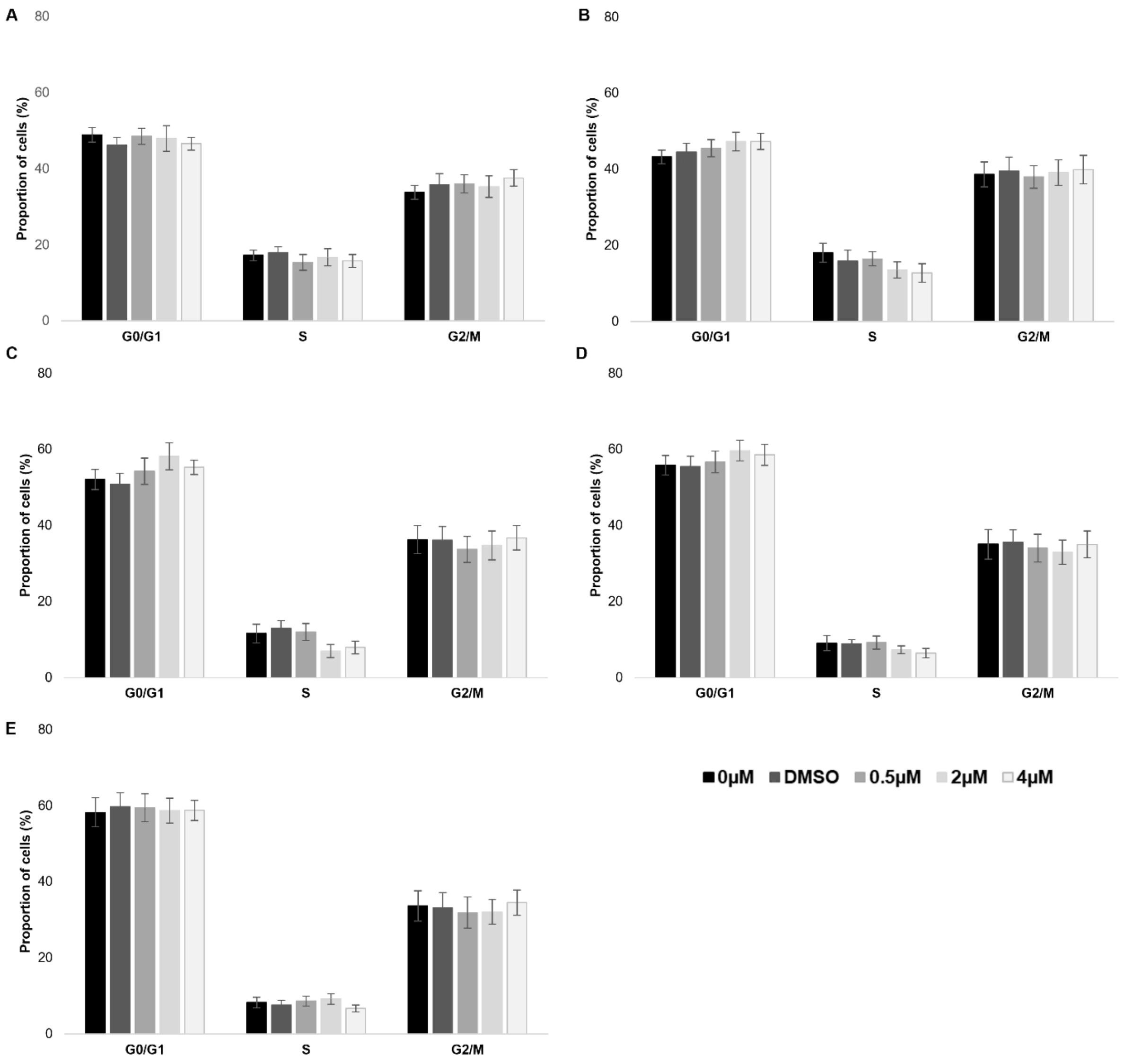


**SI. 3** Distribution of PC3 cells in the phases of the cell cycle after the treatment with the gold-NHC complex **I** at 0h (A), 24h (B), 48h (C), 72h (D), 96h (E). The results are represented as the mean of at least three independent cell cycle analysis. Statistical comparisons are made for each phase of the cell cycle. Means with dissimilar superscripts are significantly different (P < 0.05)

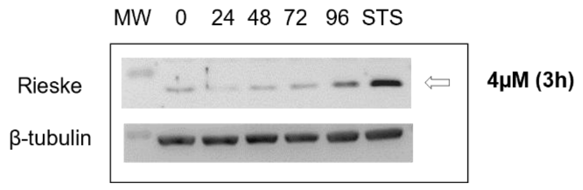


**SI. 4** Effects of the metallocarbene **I** at the concentration of 4 µM on the protein level of Rieske in PC3 cells. The compound was removed after 3 hours of treatment. Staurosporine (STS) was used as a positive control. Experiment made once
